# Biochemical mechanism of *p*-cresol removal by *Thauera aminoaromatica* S2

**DOI:** 10.64898/2026.01.30.702948

**Authors:** Oishi Sen, Pei-Hsin Wang, Prakit Saingam, Bruce J. Godfrey, Jonathan Himmelfarb, Yi Xiong, Chongle Pan, Mari Karoliina Henriikka Winkler

**Affiliations:** Department of Civil and Environmental Engineering, University of Washington; Seattle, Washington; Center for Kidney Disease Innovation, Icahn School of Medicine at Mount Sinai; School of Biological Sciences, University of Oklahoma

## Abstract

Protein-bound uremic toxins are inefficiently cleared by dialysis and contribute to complications in chronic kidney disease, motivating approaches that target their gut-derived precursors. Here we investigate anaerobic *p*-cresol metabolism by the environmental denitrifier *Thauera aminoaromatica* S2, a pathway originally evolved for aromatic pollutant degradation. Proteomic stable isotope probing with ^13^C-labeled *p*-cresol reveals strong incorporation of labeled carbon into *T. aminoaromatica* proteins, whereas parallel incubations with human fecal microbiomes show minimal incorporation, indicating limited intrinsic gut capacity for *p*-cresol utilization. Label-enriched proteins enable reconstruction of the anaerobic *p*-cresol degradation pathway and identification of key enzymes synthesized during growth on *p*-cresol. Moreover, hydrogel-encapsulated *T. aminoaromatica* remains active during co-incubation with the gut microbiome, achieving complete removal of 0.3 mM *p*-cresol in less than 10 hours, a timescale compatible with typical intestinal transit in the colon. Together, these findings establish a biochemical basis for repurposing environmental aromatic degradation pathways for gut-localized *p*-cresol removal.

## Introduction

Microbial metabolism offers powerful opportunities to address disease-relevant chemical transformations that are inaccessible to host physiology. In chronic kidney disease (CKD), this potential is particularly compelling, as several clinically relevant toxins originate from microbial metabolism in the gut yet cannot be efficiently removed once they enter systemic circulation. Genetically engineered *Escherichia coli* has been used to successfully produce human insulin (Riggs, 2021), human growth hormone (Rezaei and Zarkesh-Esfahani, 2012), human parathyroid hormone (Al-Badran and Abdul-Jabbar, 2017), somatostatin (Riggs, 2021), glucagon (Yoon et al., 2024), and non-natural higher bioalcohols such as hexanol and octanol (Sen et al., 2023), and several bacterial species have tumoricidal abilities (Duong et al., 2019; Mills et al., 2022). These examples illustrate the broader principle that non-native or engineered microbial metabolisms can be deployed for medical use.

Kidney malfunction leads to the accumulation of toxins in the blood such as urea, creatinine, and uric acid, which can be removed by dialysis (Jain et al., 2009). However, a defining and poorly addressed feature of chronic kidney disease (CKD) is the accumulation of protein-bound uremic toxins (PBUTs), including *p*-cresyl sulfate and indoxyl sulfate, which are inefficiently cleared by dialysis due to strong albumin binding (Liabeuf et al., 2011; Saingam et al., 2025). Elevated circulating levels of *p*-cresyl sulfate (PCS) and indoxyl sulfate (IS) are strongly associated with disease severity and adverse outcomes in patients with impaired kidney function (Meijers and Evenepoel, 2011). Because PBUTs originate from gut-derived metabolites and cannot be effectively removed once they enter systemic circulation, therapeutic strategies that act upstream—within the gastrointestinal tract—represent an attractive complementary approach to dialysis. Both *p*-cresol and indole contribute to PBUT accumulation; however, *p*-cresol (LD_50_ = 207 mg kg^−1^) is substantially more toxic than indole (LD_50_ = 1000 mg kg^−1^), based on oral toxicity studies in rats (SDS, Thermo Fisher Scientific). *p*-Cresol is produced by colonic microbiota through fermentation of tyrosine or phenylalanine (Zhang et al., 2025) and is subsequently sulfonated in the liver to form circulating PCS. *p*-Cresol and its derivatives exert deleterious effects on multiple organ systems, including the kidney, liver, colon, bladder, cardiovascular system, and skeletal muscle (Zhang et al., 2025). Despite successful removal of urea, uric acid, and ammonia using microencapsulated engineered *E. coli* in both *in vitro* and *in vivo* models (Jain et al., 2009), microbial strategies for eliminating PBUT precursors remain largely unexplored. This gap is notable given that current PBUT management relies primarily on dietary interventions, such as protein restriction, which can reduce circulating PCS and IS levels (Patel et al., 2012) but are difficult to sustain long term. Prebiotic and probiotic approaches aim to modulate gut microbiome composition by enriching beneficial taxa (Mitrović et al., 2023; Rossi et al., 2016), yet most focus on suppressing toxin formation rather than actively degrading PBUT precursors once formed. Consequently, the native gut microbiome retains limited intrinsic capacity to remove *p*-cresol prior to absorption.

Hydrogel-encapsulated *T. aminoaromatica* have been shown to efficiently degrade *p*-cresol under gut-relevant conditions, potentially enabling upstream removal of this uremic toxin precursor in patients with CKD (Saingam et al., 2025). This metabolic capability derives from pollutant degrading metabolic function originally evolved for aromatic compounds removal common in industrial effluents (Mechichi et al., 2002, Hsu et al., 2025), supported by the complete genome sequence of *T. aminoaromatica* (Jiang et al., 2012) and confirmed here with stable isotope probing. Notably, this strategy parallels approaches in wastewater treatment, where densely immobilized or biofilm-associated microbial communities enable higher volumetric pollutant removal rates than suspended cultures by increasing the ratio of active microorganisms to reactor volume (Winkler and van Loosdrecht, 2022). Similarly, hydrogel confinement in the gut may enhance detoxification capacity per administered volume by maintaining high local microbial activity within a compact matrix (Landreau et al., 2020).

Stable isotope probing (SIP) enables direct tracking of carbon flow through microbial metabolism by supplying substrates labeled with heavy isotopes such as ^13^C or ^14^C (Radajewski et al., 2000). Proteomic SIP extends this approach by detecting isotope incorporation into newly synthesized proteins, allowing identification of organisms and enzymes actively responsible for a given metabolic transformation (Pan et al., 2011). This strategy has been successfully applied to resolve anaerobic and aerobic degradation pathways for environmental contaminants, including identification of *Burkholderiaceae* involved in ^13^C-toluene identified (Lünsmann et al., 2016) and degradation and sufate-reducing consortia capable of degrading ^13^C-benzene (Taubert et al., 2012). Here, we apply proteomic SIP to compare *p*-cresol utilization by pure *T. aminoaromatica* S2 cultures and human fecal microbiomes by incubating both with ^13^C-labeled *p*-cresol. This approach enables reconstruction of the *p*-cresol degradation pathway in *T. aminoaromatica* while directly testing whether native gut microorganisms possess analogous metabolic capacity. Finally, we evaluated whether hydrogel-encapsulated *T. aminoaromatica* could actively remove *p*-cresol in the gut microbiome, validating our conceptual model that such therapeutic hydrogel may reduce *p*-cresol levels and alleviate kidney patient burden.

## Materials and method

### Strain and chemicals used

*T. aminoaromatica* S2 was purchased from DSMZ and was cultivated as previously described (Saingam et al., 2025). Briefly, cells were revived from glycerol stocks, plated on R2A agar, and grown in basal medium supplemented with acetate (30 mM) and nitrate (25 mM) at 30°C. Cells were harvested by centrifugation, washed twice with basal medium, and used as inocula for subsequent experiments. ^13^C-labeled *p*-cresol was purchased from Cambridge Isotope Laboratories ([*p*-Cresol (ring-^13^C_6_, 99%)-Cambridge Isotope Laboratories, CLM-7341-PK](https://isotope.com/synthetic-intermediates/p-cresol-ring-13c6-clm-7341)). All the six carbons of the benzene ring are labeled in this *p*-cresol.

### Relative toxicity of pathway intermediates compared to *p*-cresol

The toxicity of the pathway intermediates, which are not linked to Coenzyme A, relative to *p*-cresol toxicity in Fig. 3C was calculated by the formula-

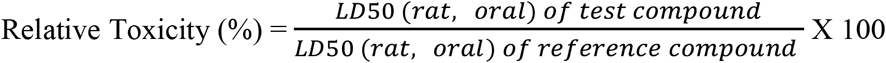

### Proteomic-SIP *Sample preparation*

Acetate-grown *T. aminoaromatica* S2 cells were harvested and incubated anaerobically in minimal medium buffered with phosphate and supplemented with 0.6 mM *p*-cresol and 10 mM KNO_3_ in Balch tubes. ^13^C-labeled *p*-cresol was supplied to Samples 1–3, while unlabeled *p*-cresol served as the control (Samples 4–6); all conditions were performed in triplicate. Human gut microbiomes were extracted from around 10 g of fecal samples by suspension in 90 ml PBS followed by low-speed centrifugation (500×g for 5 min) to remove large particulates. The supernatant was then centrifuged at 3,000 × g for 10 min to pellet microbial cells, which were resuspended and washed with 2 mL of PBS. Pellets were resuspended and incubated under identical conditions to the *T. aminoaromatica* cultures with either ^13^C-labeled or unlabeled *p*-cresol using the same experimental design. Following incubation, cells from both pure cultures and gut microbiome samples were harvested and submitted for proteomic analysis.

### Proteomic-SIP Analysis of the SIPROS output from *T. aminoaromatica* S2 proteomic samples

All contaminant and decoy proteins were removed from the SIPROS output for Samples 1–3 (^13^C-labeled) and Samples 4–6 (unlabeled controls). MS1 isotopic abundances were filtered to retain only measurements with a Q-value < 0.01 and a posterior error probability < 0.01. For each protein, MS1 isotopic abundances were averaged across all associated peptides, with a minimum requirement of three peptides per protein; proteins represented by fewer than three peptides were excluded from further analysis. The average MS1 isotopic abundance (% ^13^C incorporation) for each protein was compiled across Samples 1–6 using MS Excel. Average and standard deviations (SD) of each protein were calculated separately for Sample 1-3 (labeled group) and Sample 4-6 (control group). Proteins exhibiting substantially higher % ^13^C incorporation in the labeled group relative to controls were interpreted as being actively synthesized by *T. aminoaromatica* S2 during growth on *p*-cresol. Statistical significance was assessed using multiple t-tests implemented in GraphPad, applying the two-stage linear step-up procedure of Benjamini, Krieger, and Yekutieli with a false discovery rate (Q) of 1%. Each test was performed independently without assuming equal variance.

### Analysis of sequence homology

Protein sequences were retrieved from UniProt. Sequence homology of the FAD-linked oxidase domain protein (C4ZM78) from *T. aminoaromatica* was assessed using BLASTp against characterized *p*-cresol methylhydroxylase (PCMH) enzymes from *Geobacter metallireducens* GS-15 and *Pseudomonas putida*. In BLASTp analyses, the query refers to the *T. aminoaromatica* protein, while subjects denote reference sequences. Percent identity reflects exact amino acid matches between query and subject, whereas percent similarity includes both identical residues and conservative substitutions indicative of functional conservation.

### Analysis of the organization of genes in the genome

Operon mapper was used to detect genes belonging to a single operon. The complete genome of *T. aminoaromatica* was obtained from NCBI ([*Thauera sp*. MZ1T, complete genome - Nucleotide - NCBI](https://www.ncbi.nlm.nih.gov/nuccore/CP001281)) and genomic regions were viewed in Snapgene for analysis. Similarly, the complete genome of *G. metallireducens* GS-15 was obtained from NCBI ([Geobacter metallireducens GS-15, complete genome - Nucleotide - NCBI](https://www.ncbi.nlm.nih.gov/nuccore/CP000148)). *p*-Cresol degradative pathway genes belonging to plasmid pRA4000 of *P. putida* were obtained from NCBI ([*Pseudomonas putida* NCIMB 9866 plasmid pRA4000 p-cresol degradative pat - Nucleotide - NCBI](https://www.ncbi.nlm.nih.gov/nuccore/U96338)).

### *p*-Cresol removal by encapsulated *T. aminoaromatica* S2 in the gut microbiome Hydrogel encapsulation preparation

The cells were prepared following Saingam’s method (Saingam *et al*., 2025), also mentioned in Method 3a with an extra step to scale up acetate-grown *T. aminoaromatica* S2 up to 200 mL using a 3% (v/v) inoculum, and cultivation was repeated until the culture reached full growth. To confirm *p*-cresol removal capability, the cells were transferred to basal medium containing 0.6 mM *p*-cresol and 10 mM KNO_3_. Cells were collected slightly different from Saingam’s method: centrifugation at 1000 × g for 25 min, washed twice with basal medium, and resuspended in 1 mL of basal medium for hydrogel encapsulation. After encapsulation, 40 g of hydrogel beads were revived in 240 ml basal medium supplemented with 32 mM acetate and 10 mM KNO_3_ couple days for recovery.

### Chemical analysis

Chemical analysis followed Saingam’s method (Saingam *et al*., 2025), with modifications. Liquid samples were filtered through a 0.2 µm nylon filter (VWR, Radnor, PA) prior to measurements of *p*-cresol. *p*-Cresol concentrations were determined using an Agilent Infinity II liquid chromatography (LC) system equipped with an InfinityLab Poroshell 120 column (3.0 × 100 mm, 2.7 μm, Agilent Technologies, Part No. 683975-314T), instead of the column used in the original method. The mobile phase consisted of acetonitrile and 50 mM KH_2_PO_4_ at an approximate 1:9 (v/v) ratio, varying over the course of the run. The flow rate was 0.5 mL/min, and the column temperature was maintained at 40°C. Chromatograms were recorded at a UV wavelength of 280 nm.

### *p*-Cresol removal test

Forty grams of *T. aminoaromatica* hydrogel beads were rinsed twice with 20 mL of basal medium and then immersed in 40 mL of basal medium for 30 min for thorough washing. Subsequently, 3 g of encapsulated beads were added to 40 mL of basal medium in serum bottles. The bottles were purged with nitrogen for 15 min, sealed, and further subjected to vacuum– CO_2_ purge cycles for a total of 10 min to remove oxygen. Severn serum bottles were prepared in total: three bottles contained only *T. aminoaromatica* beads in basal medium (triplicate), three bottles contained *T. aminoaromatica* beads plus gut microbiome (triplicate), and another one serum bottle contained gut microbiome in basal medium. Gut microbiomes were processed following Method 3a with slight modifications. After centrifugation at 3,000 × g, pellets were resuspended in 10 mL PBS and centrifuged again at the same speed for 5 min. The final pellet was resuspended in 4 mL PBS, yielding approximately 5.5 mL of gut microbiome suspension. One milliliter of this suspension was added to 40 mL basal medium prepared in the same manner as above, and the remaining suspension was evenly distributed into the three serum bottles containing *T. aminoaromatica* beads. *p*-Cresol (0.3 mM) and KNO_3_ (10 mM) were added to all bottles to initiate the *p*-cresol removal test. Liquid samples were collected for measurements of *p*-cresol.

## Results and discussion

### Utilization of *p*-cresol by *T. aminoaromatica* S2

The results confirmed that *T. aminoaromatica S2* showed a very high % ^13^C incorporation in their proteins compared to proteins in *T. aminoaromatica* S2 cells fed with unlabeled *p*-cresol (Fig. 1, sample 01-06). This suggests that *p*-cresol has been degraded and utilized by *T. aminoaromatica* S2 for the synthesis of their own proteins. This result demonstrates the potential of denitrifying bacteria such as *T. aminoaromatica* as a *p*-cresol (toxin) remover from the digestive system of kidney patients. The absence of higher % ^13^C incorporation in the gut microbiome fed with ^13^C labeled *p*-cresol (Fig. 1, Samples 07-09) compared to gut microbiome fed with unlabeled *p*-cresol (Fig. 1, Samples 10-12), shows that the human gut microbiome lacks an efficient mechanism of removing *p*-cresol on its own, which eventually is absorbed and accumulated in kidney patients leading to adverse health effects.

**Figure 1:**
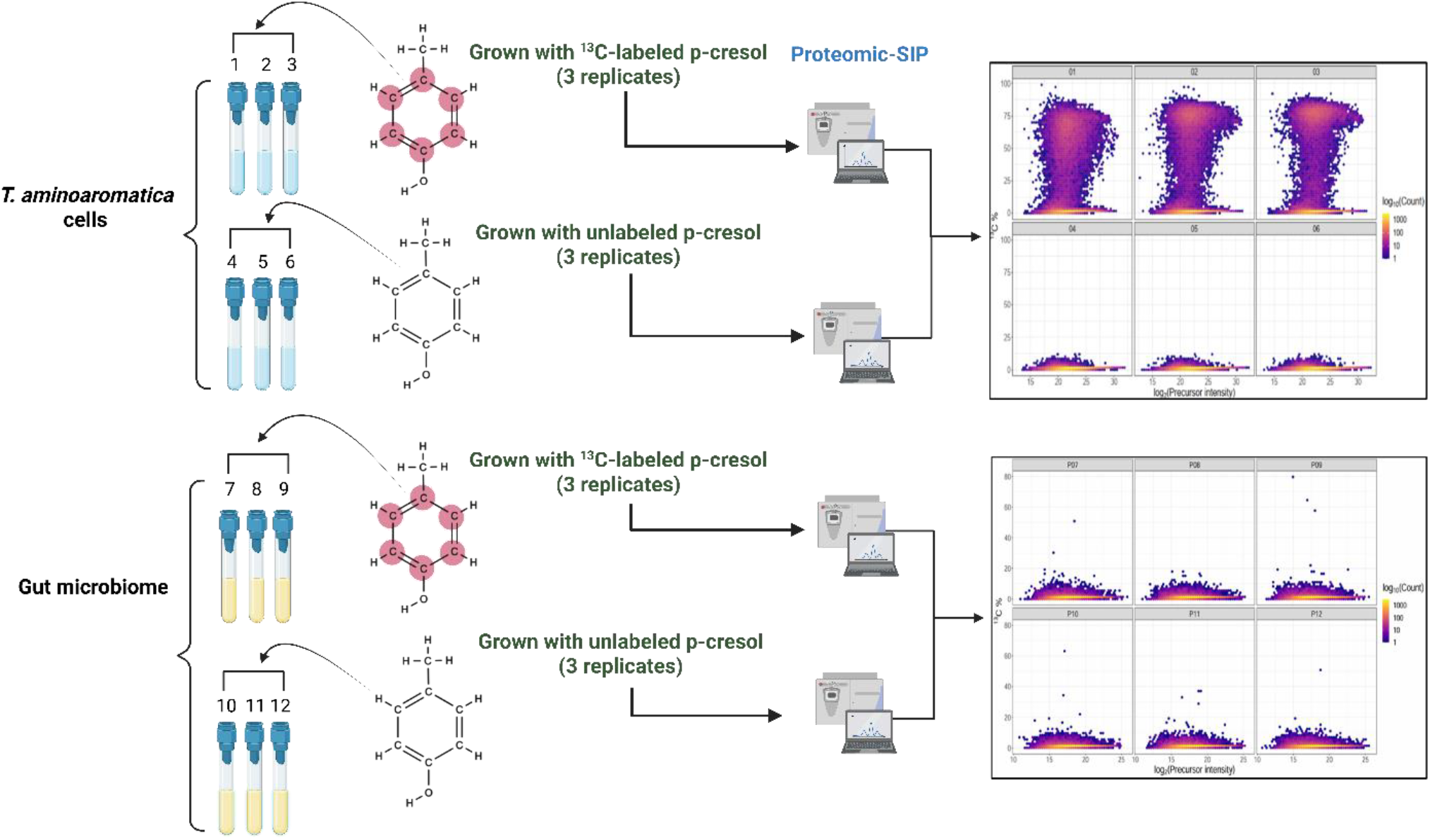
Proteomic stable isotope probing reveals *p*-cresol assimilation by *T. aminoaromatica* but not by the native gut microbiome. Planktonic *T. aminoaromatica* cultures incubated with ^13^C-labeled *p*-cresol (01–03) showed substantial incorporation of ^13^C into cellular proteins compared with unlabeled controls (04–06), indicating active *p*-cresol degradation and assimilation. In contrast, human gut microbiome samples incubated with ^13^C-labeled *p*-cresol (P07–09) showed no detectable increase in ^13^C incorporation relative to unlabeled controls (P10–12), indicating an absence of robust *p*-cresol utilization under these conditions. Pink denotes ^13^C-labeled carbon atoms derived from *p*-cresol.

### *T. aminoaromatica* utilizes a specific pathway for anaerobic degradation of *p*-cresol

It is well established that anaerobic degradation of aromatic hydrocarbons, including *p*-cresol, proceeds via the central intermediate benzoyl-CoA (Harwood et al., 1998). Using proteomic stable isotope probing (SIP), we resolved the enzymatic pathway by which *T. aminoaromatica* anaerobically degrades and assimilates *p*-cresol (Fig. 2A). Substantial ^13^C incorporation was detected across nearly all pathway enzymes (Fig. 2B), indicating active synthesis during growth on labeled substrate. With the exception of 3-hydroxyacyl-CoA dehydrogenase (C4ZJ65) and acetyl-CoA C-acyltransferase (C4ZJ66), these proteins ranked among the top 40 (top 7.5%) of the 533 detected proteins based on maximum ^13^C incorporation (Table 1). This enrichment pattern demonstrates that carbon derived from *p*-cresol is directly assimilated into newly synthesized enzymes within the anaerobic benzoyl-CoA degradation pathway.

**Table 1:** Top 40 proteins showing the highest % ^13^C incorporation.

| Uniprot Protein ID | Gene name | Gene locus | Description | % <sup>13</sup>C incorporation (labeled - unlabeled) |
| --- | --- | --- | --- | --- |
| C4ZL35 | - | Tmz1t_0198 | 4Fe-4S dicluster domain-containing protein | 68.18125 |
| C4ZL36 | - | Tmz1t_0199 | Molybdopterin oxidoreductase | 70.44729 |
| C4ZL39 | - | Tmz1t_0202 | XdhC family protein | 68.79734 |
| C4ZMH5 | *hcrC* | Tmz1t_0224 | 4-Hydroxybenzoyl-CoA reductase subunit gamma | 55.33344 |
| C4ZMH6 | *hcrA* | Tmz1t_0225 | 4-Hydroxybenzoyl-CoA reductase subunit alpha | 59.59719 |
| C4ZMH7 | *hcrB* | Tmz1t_0226 | 4-Hydroxybenzoyl-CoA reductase subunit beta | 56.5916 |
| C4ZIR1 | - | Tmz1t_0419 | Glutaryl-CoA dehydrogenase (ETF) | 62.04005 |
| C4ZK76 | - | Tmz1t_1165 | Heat shock protein Hsp20 | 39.46054 |
| C4ZLY8 | - | Tmz1t_1284 | Universal stress protein | 37.42214 |
| C4ZP71 | *paaF* | Tmz1t_1509 | Phenylacetate-coenzyme A ligase | 40.97275 |
| C4ZM76 | - | Tmz1t_1838 | Phenol metabolism protein | 42.10725 |
| C4ZM77 | - | Tmz1t_1839 | Aldehyde Dehydrogenase | 40.6361 |
| C4ZM78 | - | Tmz1t_1840 | FAD linked oxidase domain protein | 43.48229 |
| C4KAW3 | - | Tmz1t_2940 | ABC transporter ATP-binding protein | 57.63781 |
| C4KAW6 | - | Tmz1t_2943 | ABC transporter substrate-binding protein | 52.57462 |
| C4KAW8 | - | Tmz1t_2945 | GCN5-related N-acetyltransferase | 55.71943 |
| C4KAW9 | - | Tmz1t_2946 | GCN5-related N-acetyltransferase | 51.57831 |
| C4KAX1 | *bcrD* | Tmz1t_2948 | Benzoyl-CoA reductase subunit D | 55.59783 |
| C4KAX2 | *bcrA* | Tmz1t_2949 | Benzoyl-CoA reductase subunit A | 57.72399 |
| C4KAX3 | *bcrB* | Tmz1t_2950 | Benzoyl-CoA reductase subunit B | 56.1617 |
| C4KAX4 | *bcrC* | Tmz1t_2951 | Benzoyl-CoA reductase subunit C | 56.16284 |
| C4KAX5 | *dch* | Tmz1t_2952 | Cyclohexa-1,5-dienecarbonyl-CoA hydratase | 60.466 |
| C4KAX6 | *oah* | Tmz1t_2953 | 6-Oxocyclohex-1-ene-1-carbonyl-CoA hydratase | 58.34701 |
| C4KAX7 | *had* | Tmz1t_2954 | 6-Hydroxycyclohex-1-ene-1-carbonyl-CoA dehydrogenase | 57.55355 |
| C4KAX8 | - | Tmz1t_2955 | 2-Oxoacid:acceptor oxidoreductase subunit alpha | 52.51509 |
| C4KAX9 | - | Tmz1t_2956 | 2-Oxoacid:ferredoxin oxidoreductase subunit beta | 52.90122 |
| C4KAY1 | - | Tmz1t_2958 | CBS domain-containing protein | 44.62154 |
| C4KAY2 | - | Tmz1t_2959 | Molybdopterin oxidoreductase | 68.41679 |
| C4KAY5 | *padE* | Tmz1t_2962 | NADH-dependent phenylglyoxylate dehydrogenase subunit gamma | 68.86667 |
| C4KAY7 | *padG* | Tmz1t_2964 | NADH-dependent phenylglyoxylate dehydrogenase subunit alpha | 68.54613 |
| C4KAY8 | *padH* | Tmz1t_2965 | NADH-dependent phenylglyoxylate dehydrogenase subunit epsilon | 65.44156 |
| C4KAY9 | *padI* | Tmz1t_2966 | NADH-dependent phenylglyoxylate dehydrogenase subunit beta | 68.62844 |
| C4KAZ0 | *paaK* | Tmz1t_2967 | Phenylacetate-coenzyme A ligase | 68.3761 |
| C4KAZ1 | - | Tmz1t_2968 | Organic acid-CoA ligase (ADP forming) | 67.96152 |
| C4KB11 | - | Tmz1t_2988 | Nitrogen-fixing NifU domain protein | 64.68399 |
| C4KB23 | - | Tmz1t_3000 | AMP-dependent synthetase and ligase | 63.13367 |
| C4KBF2 | - | Tmz1t_3131 | ABC transporter related | 54.82792 |
| C4KBF3 | - | Tmz1t_3132 | ABC transporter ATP-binding protein | 62.53935 |
| C4KBF7 | - | Tmz1t_3136 | Benzoate-CoA ligase family | 51.94367 |

**Figure 2:**
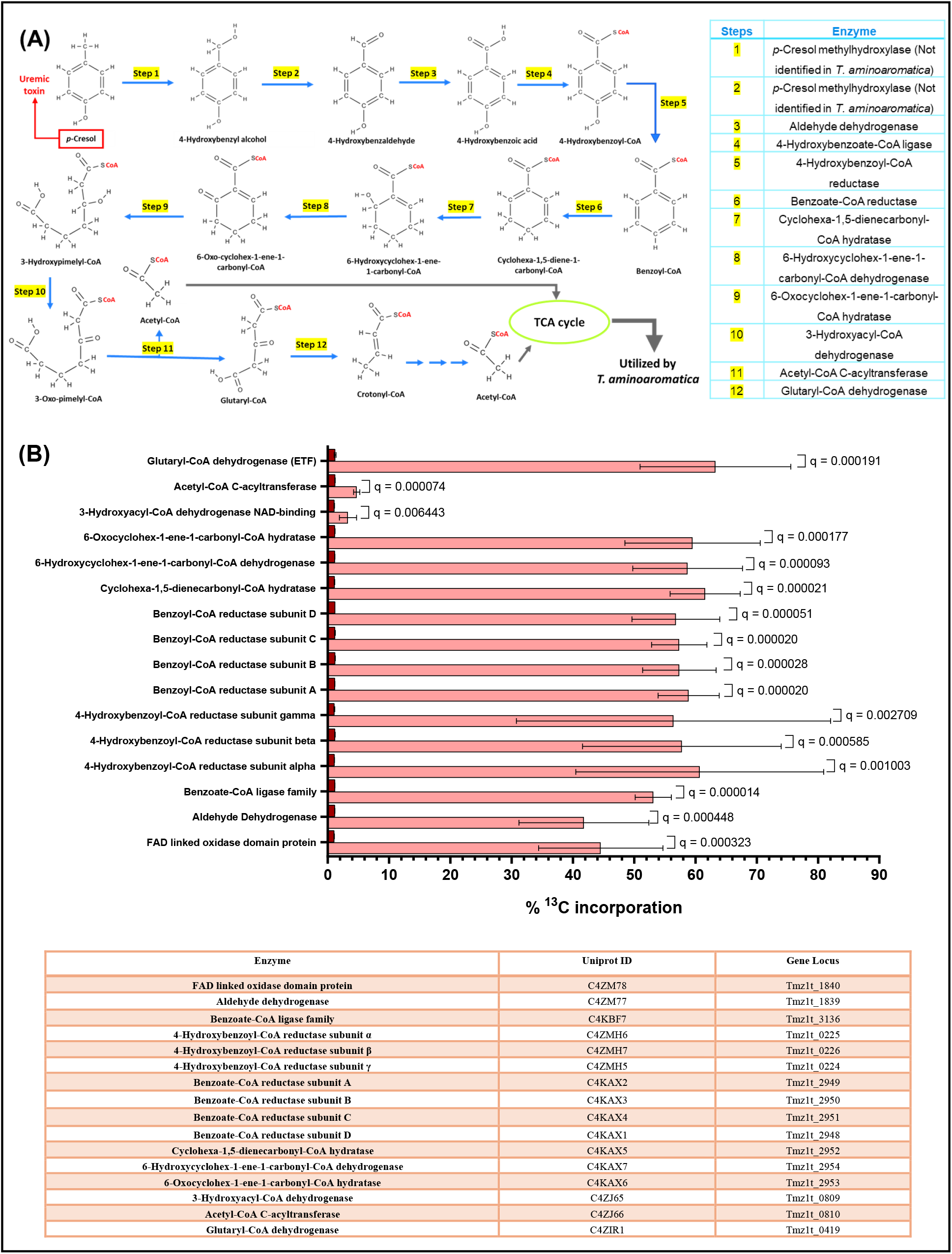
Anaerobic *p*-cresol degradation pathway resolved in *T. aminoaromatica*. (A) Reconstructed anaerobic pathway converting *p*-cresol to acetyl-CoA in *T. aminoaromatica*. Enzymes catalyzing steps 3–12 were identified; enzymes responsible for steps 1–2 remain unresolved. (B) ^13^C incorporation into enzymes of the *p*-cresol degradation pathway following growth on ^13^C-labeled *p*-cresol (pink bars) compared to growth on unlabeled *p*-cresol (brown bars). Bars represent mean ± SD (n = 3). Significance assessed using multiple testing with false discovery rate correction (q < 0.05).

### FAD linked oxidase domain protein (C4ZM78) shows significant homology with known PCMH

*p*-Cresol methylhydroxylase (PCMH) catalyzes the initial oxidation of *p*-cresol to 4-hydroxybenzyl alcohol and subsequently to 4-hydroxybenzaldehyde in several anaerobic and facultative anaerobic bacteria (Jõesaar et al., 2010; Peters et al., 2007). PCMH is a two-component enzyme composed of a FAD-binding flavoprotein subunit harboring the catalytic active site and a c-type cytochrome subunit that mediates electron transfer (Jõesaar et al., 2010). In *Pseudomonas putida*, the equivalent activity is annotated as 4-cresol dehydrogenase, whereas the corresponding enzyme has not been previously assigned in *T. aminoaromatica*.

To identify the enzyme responsible for the conversion of *p*-cresol to 4-hydroxybenzaldehyde in *T. aminoaromatica*, we combined proteomic stable isotope probing with comparative sequence and gene neighborhood analyses. In the proteomic-SIP dataset, a protein annotated as an FAD-linked oxidase domain protein (Uniprot C4ZM78; 516 aa) exhibited consistently high ^13^C incorporation during growth on labeled *p*-cresol (Fig. 2B), indicating active synthesis and direct assimilation of *p*-cresol–derived carbon. Uniprot annotation further indicates that C4ZM78 contains a PCMH-type FAD-binding domain.

Pairwise BLASTp analysis revealed strong homology between C4ZM78 and well-characterized PCMH enzymes, including the flavoprotein subunit of 4-cresol dehydrogenase from *P. putida* (P09788) and the alpha-prime subunit of PCMH from *Geobacter metallireducens* (Q39TS0). Alignments covered ≥ 98% of both query and subject sequences, with sequence similarities of 75.4% (*P. putida*) and 69.3% (*G. metallireducens*), and identities of 58.8% and 52.4%, respectively. In both cases, E-values were zero and bit scores exceeded 570, indicating robust evolutionary conservation unlikely to arise by chance (Fig. 3A).

**Figure 3:**
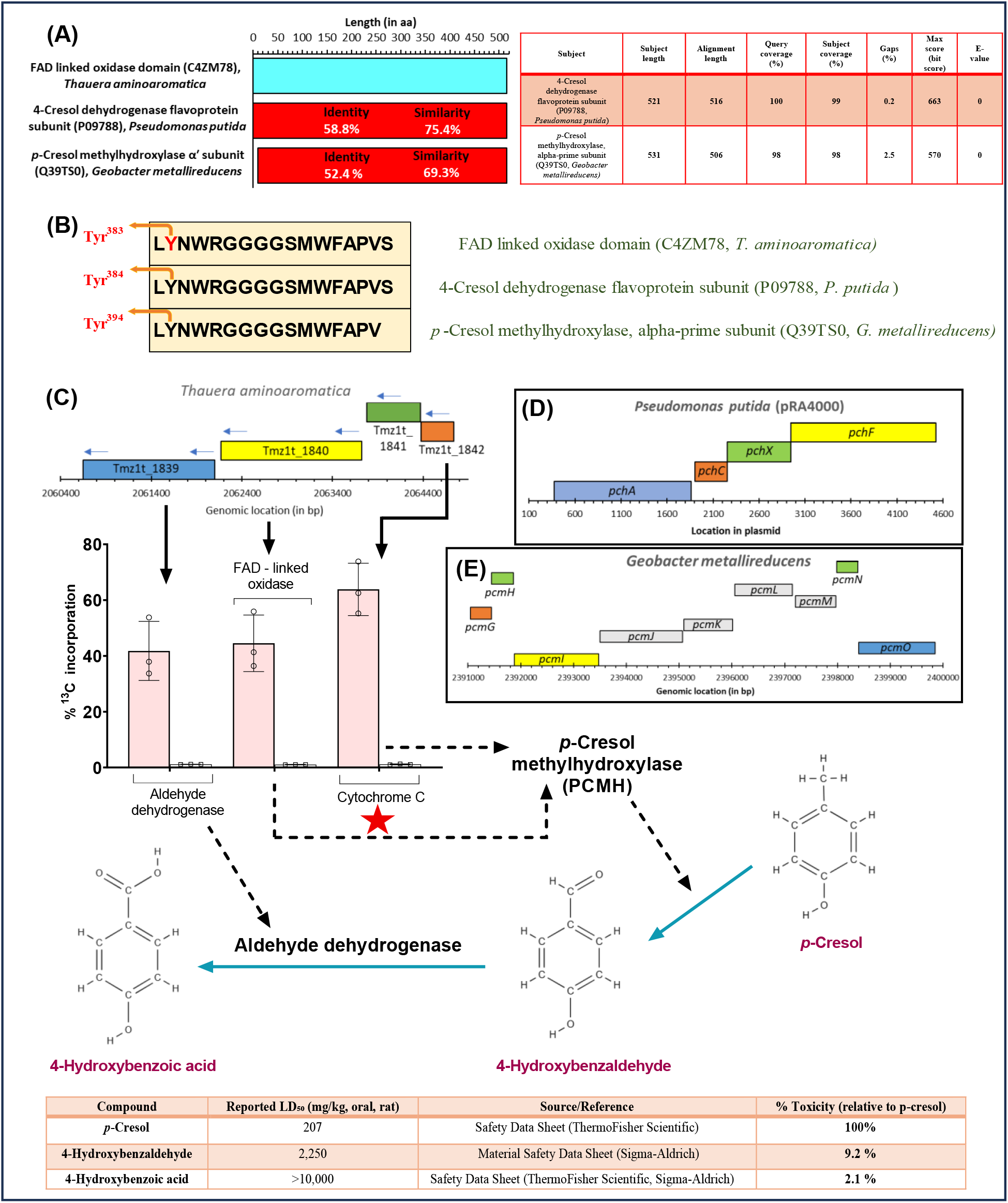
Conservation of *p*-cresol methylhydroxylase (PCMH) architecture and gene organization across *p*-cresol–degrading bacteria. (A) Pairwise sequence alignment of the FAD-linked oxidase domain protein from *T. aminoaromatica* (C4ZM78) with characterized PCMH enzymes using BLASTp, showing percent identity, similarity, and query coverage. (B) Conservation of the FAD-binding active-site tyrosine residue and surrounding amino acid motif across *T. aminoaromatica, P. putida*, and *G. metallireducens*. (C) Operon structure in *T. aminoaromatica* encoding the FAD-linked oxidase subunit (C4ZM78), associated cytochrome subunit, and aldehyde dehydrogenase, with corresponding gene loci and catalyzed reactions indicated. (D) Gene neighborhood surrounding the 4-cresol dehydrogenase complex encoded on plasmid pRA4000 in *P. putida*. E) Neighbouring genes adjacent to genes encoding *p*-cresol methylhydroxylase subunits in *G. metallireducens*. In all the bacteria shown here, there is presence of genes close to each other that encodes for i) a cytochrome subunit (orange), ii) a FAD subunit (yellow) of the enzyme catalysing the conversion of *p*-cresol to 4-hydroxybenzaldehyde, iii) the enzyme catalysing the conversion of 4-hydroxybenzaldehyde → 4-hydroxybenzoic acid (blue).

In *P. putida*, Tyr^384^ of the flavoprotein subunit (*pchF*) forms a covalent linkage with FAD, which is essential for catalytic activity (Efimov et al., 2001). Similarly, in *G. metallireducens*, Tyr^394^ of the PCMH alpha-prime subunit (*pcmI*) serves as the active-site residue for covalent FAD attachment, and deletion of *pcmI* abolishes *p*-cresol degradation (Johannes et al., 2008; Chaurasia et al., 2015). Both residues are embedded within a conserved NWRGGGGSMWFAPV motif and preceded by a conserved leucine residue, features thought to stabilize the catalytic environment (Fig. 3B).

Notably, C4ZM78 from *T. aminoaromatica* contains an identically positioned tyrosine residue (Tyr^383^) flanked by the same conserved neighboring amino acids, strongly suggesting that this residue serves as the catalytic FAD-binding site. Together, the proteomic-SIP evidence, sequence homology, conserved active-site architecture, and gene neighborhood organization support the assignment of C4ZM78 as the PCMH-like flavoprotein subunit mediating the initial oxidation of *p*-cresol in *T. aminoaromatica*.

### Conserved gene neighbourhood or synteny observed in the genes encoding enzymes facilitating the conversion of *p*-cresol to 4-hydroxybenzoic acid in *T. aminoaromatica, P. putida and G. metallireducens*

In *T. aminoaromatica*, the gene encoding the FAD-linked oxidase subunit (C4ZM78) is located within an operon spanning Tmz1t_1839–1842, which encodes a cytochrome *c* subunit (C4ZM80), a hypothetical protein of unknown function (C4ZM79), the FAD-linked oxidase (C4ZM78), and an aldehyde dehydrogenase (C4ZM77) that catalyzes the conversion of 4-hydroxybenzaldehyde to 4-hydroxybenzoic acid (Fig. 3C). This operon architecture closely parallels the organization of *p*-cresol degradation genes in other well-characterized bacteria. In *P. putida*, the pRA4000 plasmid encodes the corresponding enzymatic functions in the order *pchA–pchC–pchX–pchF*, enabling conversion of *p*-cresol to 4-hydroxybenzaldehyde (Fig. 3D). Similarly, *G. metallireducens* harbors a conserved operon (*pcmG–O*) encoding cytochrome, FAD-binding, and aldehyde dehydrogenase components required for *p*-cresol methylhydroxylase activity (Fig. 3E). Notably, *T. aminoaromatica* contains adjacent genes encoding the FAD-linked oxidase, cytochrome *c*, and aldehyde dehydrogenase within a single operon, mirroring the conserved gene synteny observed in *P. putida* and *G. metallireducens*. Together, these genes encode the enzymatic machinery required for the conversion of *p*-cresol to 4-hydroxybenzoic acid. Consistent with this pathway, accumulation of 4-hydroxybenzoic acid during growth of *T. aminoaromatica* S2 on *p*-cresol has been reported previously (Saingam *et al*., 2025).

Proteomic stable isotope probing further supports this route: high ^13^C incorporation was observed in both the FAD-linked oxidase subunit (C4ZM78) and the aldehyde dehydrogenase (C4ZM77), indicating active synthesis during growth on labeled *p*-cresol. Although the cytochrome *c* subunit (C4ZM80) is small (~100 aa) and detected by a single peptide—below the threshold for confident protein-level quantification—that peptide consistently exhibited high ^13^C incorporation across replicates (Fig. 3C), supporting its functional involvement. Importantly, the primary intermediates generated by this pathway—4-hydroxybenzaldehyde and 4-hydroxybenzoic acid—are substantially less toxic than *p*-cresol based on reported oral LD_50_ values in rats. Thus, even transient release of these intermediates would be expected to pose lower toxic risk than continued accumulation of *p*-cresol, further supporting the detoxifying nature of this metabolic route.

### Subsequent steps in the anaerobic degradation of *p*-cresol in *T. aminoaromatica*

In the current study, high % ^13^C incorporation was seen in benzoate-CoA ligase family protein when *T. aminoaromatica* S2 was grown on labeled *p*-cresol, indicating active synthesis during *p*-cresol metabolism. In *Thauera aromatica*, benzoate-CoA ligase coded by *bclA* primarily activates benzoic acid to benzoyl-CoA but can also accept some other aromatic substrates. *T. aromatica* also encodes a distinct 3-hydroxybenzoate-CoA/4-hydroxybenzoate-CoA ligase that converts 4-hydroxybenzoic acid to 4-hydroxybenzoyl-CoA. In contrast, *Rhodopseudomonas palustris* encodes a 4-hydroxybenzoate-CoA/benzoate-CoA ligase (HbaA) that accepts both benzoate and 4-hydroxybenzoate as its substrate (Uniprot). Whether the benzoate-CoA ligase enzyme induced in *T. aminoaromatica*, which is actively synthesized in the presence of *p*-cresol in the current study, can accept 4-hydroxybenzoic acid remains unknown. Nevertheless, the induction of 4-hydroxybenzoyl-CoA reductase during growth on *p*-cresol suggests that the *p*-cresol degradation pathway proceeds via the formation of 4-hydroxybenzoyl-CoA. Significantly high % ^13^C incorporation in the α, β and γ subunits of 4-hydroxybenzoyl-CoA reductase was observed (Fig. 2B), suggesting the active synthesis of the enzyme and the consequent conversion of 4-hydroxybenzoyl-CoA to benzoyl-CoA (Fig. 2A, Step 5; Fig. 4A). The genes encoding the α, β, and γ subunits (*hcrA–C*) are organized in a single operon in *T. aminoaromatica* (Fig. 4A, left), which is likely induced in the presence of *p*-cresol.

**Figure 4:**
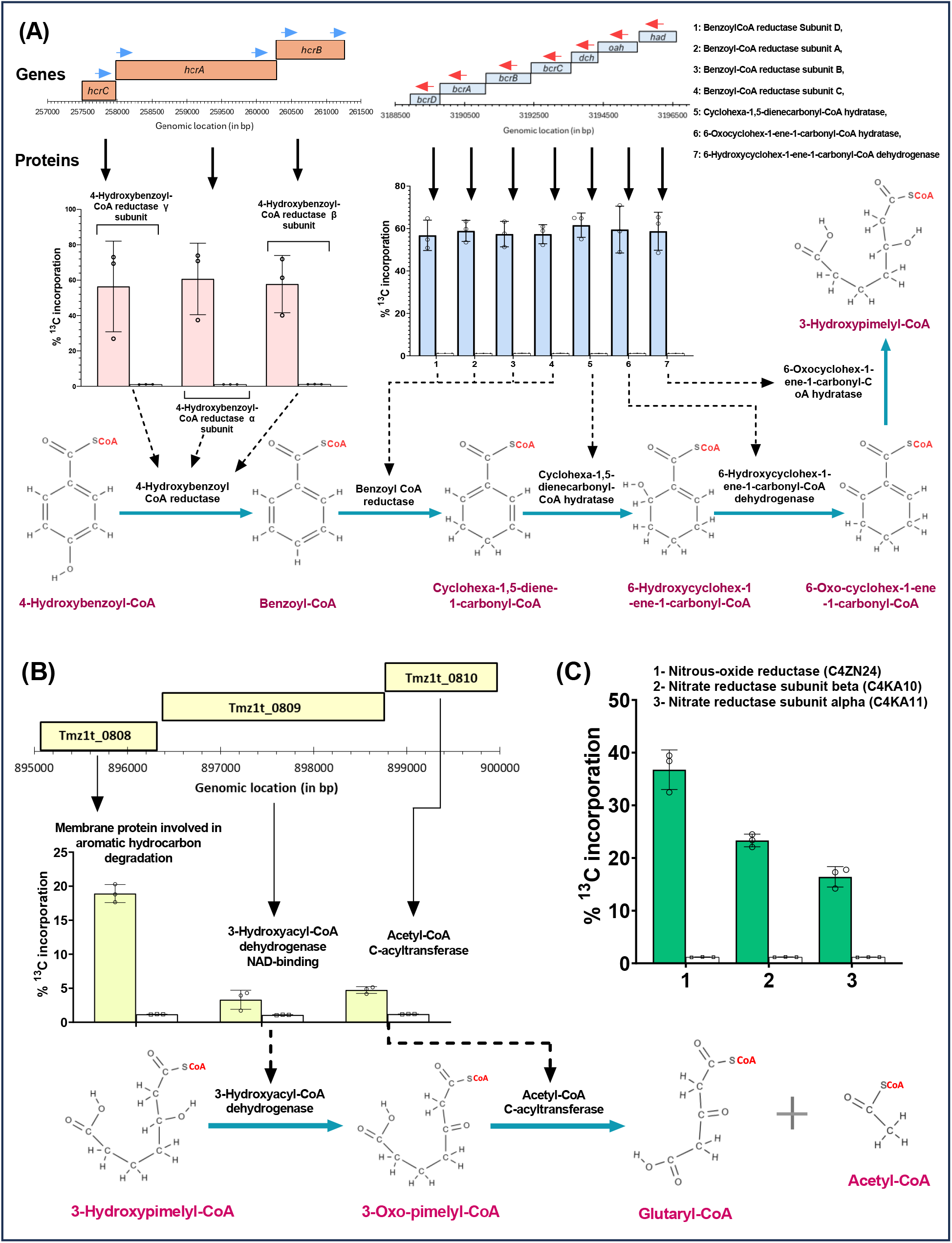
Operon organization and denitrification activity during anaerobic *p*-cresol metabolism. (A) Operon structure and encoded enzymes mediating conversion of 4-hydroxybenzoyl-CoA to benzoyl-CoA (*hrcA–C*, pink) and subsequent benzoyl-CoA degradation (*bcrA–D, dch, oah, had*; blue). (B) Operon encoding enzymes converting 3-hydroxypimelyl-CoA to glutaryl-CoA and acetyl-CoA. (C) ^13^C incorporation into proteins involved in denitrification during growth on ^13^C-labeled *p*-cresol, indicating active coupling of aromatic degradation to nitrate respiration. Error bars represent SD (n = 3).

In *T. aminoaromatica*, anaerobic benzoyl-CoA degradation proceeds via stepwise conversion to crotonyl-CoA (Fig. 2A, steps 6–12) through a series of well-defined intermediates, including cyclohexa-1,5-diene-1-carbonyl-CoA, 6-hydroxycyclohex-1-ene-1-carbonyl-CoA, 6-oxo-cyclohex-1-ene-1-carbonyl-CoA, 3-hydroxypimelyl-CoA, 3-oxo-pimelyl-CoA, and glutaryl-CoA. Enzymes catalyzing each step in this pathway exhibited significantly elevated ^13^C incorporation (Fig. 2B), consistent with active synthesis and assimilation of carbon derived from the labelled substrate. This pathway corresponds to the canonical anaerobic benzoyl-CoA degradation route described in other denitrifying bacteria (Wischgoll et al., 2009; Carmona et al., 2009). In *T. aminoaromatica*, the genes encoding the enzymes responsible for conversion of benzoyl-CoA to 3-hydroxypimelyl-CoA (*bcrA–D, dch, oah, had*) are co-localized within a single operon (Fig. 4A, right), supporting coordinated regulation of this central catabolic module.

In the anaerobic benzoyl-CoA degradation pathway, 3-hydroxypimelyl-CoA is oxidized to 3-oxo-pimelyl-CoA by 3-hydroxyacyl-CoA dehydrogenase, followed by thiolytic cleavage to glutaryl-CoA by acetyl-CoA C-acyltransferase, with concomitant release of acetyl-CoA (Peters et al., 2007; Carmona et al., 2009). The liberated acetyl-CoA can subsequently enter central metabolism via the tricarboxylic acid (TCA) cycle. In *T. aminoaromatica*, the genes encoding 3-hydroxyacyl-CoA dehydrogenase (C4ZJ65; EC 1.1.1.35) and acetyl-CoA C-acyltransferase (C4ZJ66; EC 2.3.1.16) are co-localized within a single operon together with a third gene (C4ZJ64) annotated as a membrane protein involved in aromatic hydrocarbon degradation (UniProt; Fig. 4B). All three proteins exhibited high ^13^C incorporation relative to unlabeled controls, consistent with their active synthesis during growth on labeled *p*-cresol.

Glutaryl-CoA is subsequently converted to crotonyl-CoA by glutaryl-CoA dehydrogenase (C4ZIR1; Fig. 2A, step 12), which also showed significantly elevated ^13^C incorporation (Fig. 2B). Crotonyl-CoA is further metabolized through β-oxidation–like reactions to acetyl-CoA, completing the assimilation of *p*-cresol–derived carbon into central metabolism. Collectively, these data demonstrate that *T. aminoaromatica* anaerobically degrades *p*-cresol and incorporates its carbon into newly synthesized biomass.

As a denitrifying bacterium, *T. aminoaromatica* couples this metabolic process to nitrate respiration, in which nitrate is sequentially reduced to dinitrogen (NO_3_^−^ → NO_2_^−^ → NO → N_2_O → N_2_) (Zumft, 1997; Bedmar et al., 2005). Energy conservation under anoxic conditions is mediated by nitrate, nitrite, nitric oxide, and nitrous oxide reductases (Knowles, 1982; Lecomte et al., 2018). In this study, nitrous oxide reductase as well as the α and β subunits of nitrate reductase exhibited significantly elevated ^13^C incorporation (Fig. 4C), indicating coordinated induction of denitrification and *p*-cresol catabolism during growth on the labeled substrate.

### Other proteins showing significantly high % ^13^C incorporation (top 40 proteins)

Phenylacetate is known to be converted to the central intermediate benzoyl-CoA via sequential conversion to phenylacetyl-CoA and phenylglyoxalate (Hirsch et al., 1998; Harwood et al., 1998). Phenylacetate is activated to phenylacetyl-CoA by phenylacetate-CoA ligase (EC 6.2.1.21, ENZYME - 6.2.1.30 phenylacetate--CoA ligase). Phenylglyoxalate is then converted to benzoyl-CoA by phenylglyoxylate dehydrogenase (EC 1.2.1.58, ENZYME - 1.2.1.58 phenylglyoxylate dehydrogenase (acylating)), a multienzyme complex composed of 5 subunits - α, β, γ, δ and ε (Hirsch et al., 1998). In *T. aminoaromatica*, the genes *padE–I* and *paaK* (Tmz1t_2962–2967) form a single operon. According to NCBI annotation, *padE–I* encode the α–ε subunits of phenylglyoxylate dehydrogenase, while *paaK* encodes phenylacetate-CoA ligase. In the present dataset, the PadF subunit (C4KAY6) was not detected, likely due to its small size (~100 aa), which falls below reliable proteomic detection thresholds. In contrast, proteins encoded by *padE* (C4KAY5), *padG–I* (C4KAY7–9), and *paaK* (C4KAZ0) exhibited high ^13^C incorporation, indicating active synthesis of enzymes associated with phenylacetate metabolism. This pattern suggests that growth on *p*-cresol may nonspecifically induce overlapping aromatic degradation pathways, consistent with shared structural features and regulatory cross-talk between *p*-cresol and phenylacetate metabolism.

Across the proteome, the 40 proteins exhibiting the highest ^13^C incorporation among 533 detected proteins showed net labeling values ranging from ~37.4% to 70.4% (Table 1). Many of these proteins lack detailed functional annotation in NCBI or UniProt, highlighting gaps in current pathway characterization. Notably, proteins involved in redox and transport functions—including molybdopterin oxidoreductases, ABC transporter ATP-binding proteins, GCN5-related N-acetyltransferases, and the α and β subunits of 2-oxoacid:acceptor oxidoreductase—were actively synthesized during growth on *p*-cresol. In Table 1, gene loci are ordered by genomic position, with color coding used to denote operon membership, facilitating interpretation of coordinated pathway induction.

### Encapsulated *T. aminoaromatica* S2 showed *p*-cresol removal in gut microbiome

Proteomic analyses showed that *T. aminoaromatica* S2 expresses proteins involved in *p*-cresol utilization in the basal medium, whereas the gut microbiome exhibited no detectable proteins associated with *p*-cresol degradation. Consistent with this, our previous work demonstrated that hydrogel-encapsulated *T. aminoaromatica* S2 enhanced *p*-cresol removal efficiency compared to planktonic cultures (Saingam et al., 2025). Building on these findings, we evaluated encapsulated *T. aminoaromatica S2* as a gut compatible *p*-cresol sink capable of functioning in presence of the native gut microbiome.

Fig. 5 compares *p*-cresol removal by hydrogel encapsulated *T. aminoaromatica* S2 alone (blue, results triplicate) and in co-incubation with gut microbiota (orange, results triplicate) over 50 hours, approximating colonic transit time. The gut microbiome alone showed no detectable *p*-cresol removal. Encapsulated *T. aminoaromatica* S2 beads retained robust activity in the presence of the gut microbiome, achieving complete *p*-cresol less than 10 hours and supporting its potential for gut-localized *p*-cresol mitigation.

**Figure 5:**
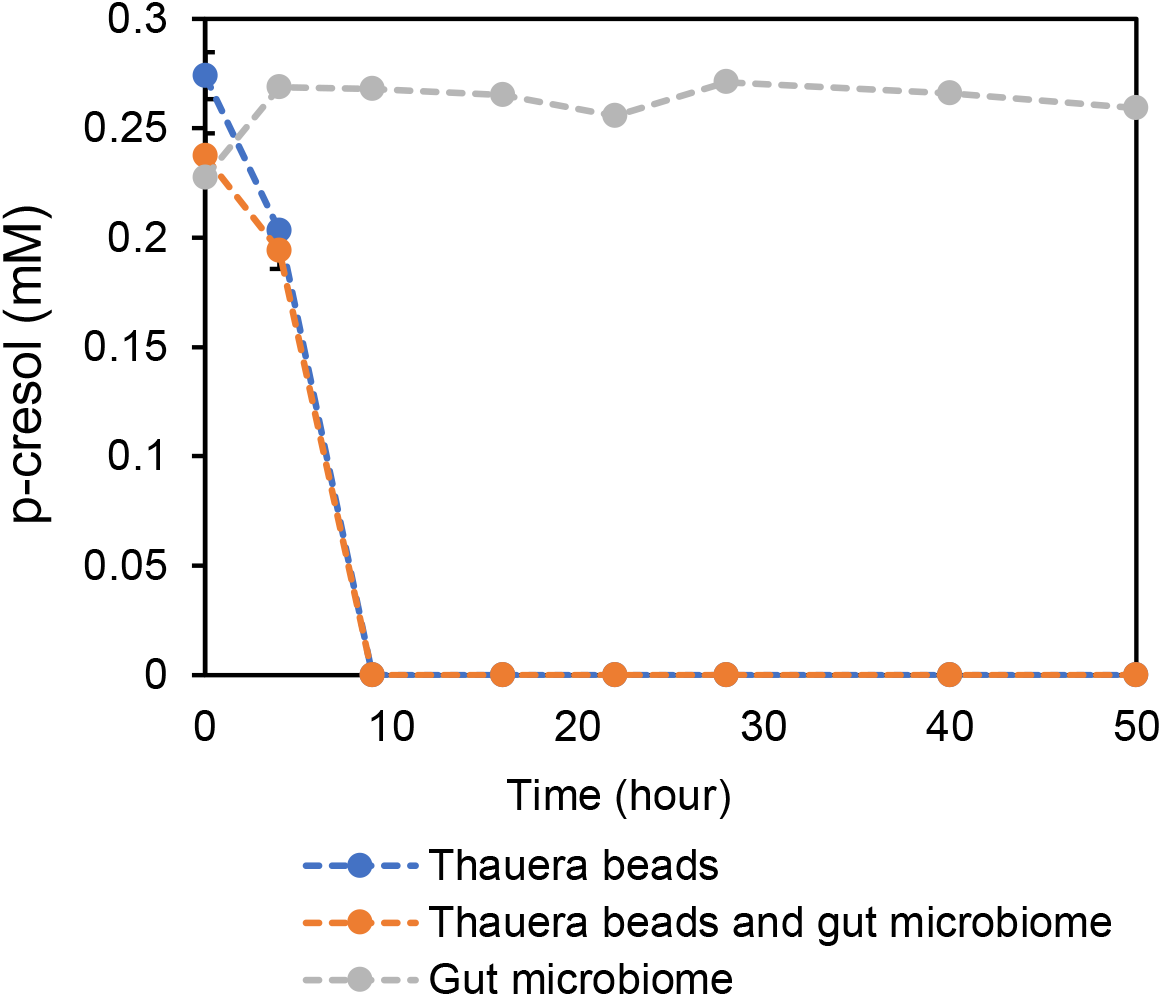
*p*-Cresol removal by *T. aminoaromatic* S2 beads. With the addition of *T. aminoaromatic* S2 in the gut microbiome environment, *p*-cresol was removed.

## Conclusions

This study identifies and functionally resolves an anaerobic *p*-cresol degradation pathway in *T. aminoaromatica*, a metabolic capability originally evolved for degradation of aromatic compounds in anoxic sludge (Mechichi et al., 2002) and largely absent from the native human gut microbiome. When densely confined within hydrogel matrices, this pathway enables effective *p*-cresol removal under gut-relevant anoxic conditions, consistent with our prior demonstrations of hydrogel-encapsulated *T. aminoaromatica* functioning in distal colon–like conditions (Saingam et al., 2025). Proteomic analyses further show that *T. aminoaromatica* actively expresses *p*-cresol utilization proteins, whereas no comparable machinery was detected in gut microbiome incubations, supporting the limited capacity of the native community to remove this precursor. Importantly, encapsulated *T. aminoaromatica* retained strong *p*-cresol removal activity even in the presence of the gut microbiome, underscoring the effectiveness of hydrogel confinement as a strategy for introducing non-native detoxification functions into the gut lumen. Proteomic-SIP–based pathway reconstruction revealed that *p*-cresol is converted into intermediates such as 4-hydroxybenzaldehyde and 4-hydroxybenzoic acid, which exhibit substantially lower toxicity than *p*-cresol. Together, these features suggest that this pathway inherently reduces toxic risk while effectively lowering *p*-cresol burden.

Overall, this work establishes a wastewater-inspired (Winkler and Straka, 2019), hydrogel-enabled strategy (Gottshall et al., 2021; Li et al., 2023) confining non-native metabolic pathways to the gut lumen, thereby unlocking new capacity for upstream protein-bound uremic toxin mitigation. More broadly, it highlights a generalizable framework for translating environmental biocatalysis into gut-targeted therapeutic concepts for chronic kidney disease and related metabolic disorders.

## Acknowledgments

We thank Britt Abrahamson and Zach Flinkstrom for their assistance with Protein Information Pipeline processing and for valuable discussions. This project was funded in part by NIH 1R01DK130815-01 from NIDDK as well as R01AT011618 from NCCIH and NIGMS.

